# Inference of the distribution of selection coefficients for new nonsynonymous mutations using large samples

**DOI:** 10.1101/071431

**Authors:** Bernard Y. Kim, Christian D. Huber, Kirk E. Lohmueller

## Abstract

The distribution of fitness effects (DFE) has considerable importance in population genetics. To date, estimates of the DFE come from studies using a small number of individuals. Thus, estimates of the proportion of moderately to strongly deleterious new mutations may be unreliable because such variants are unlikely to be segregating in the data. Additionally, the true functional form of the DFE is unknown, and estimates of the DFE differ significantly between studies. Here we present a flexible and computationally tractable method, called Fit∂a∂i, to estimate the DFE using the site frequency spectrum from a large number of individuals. We apply our approach to the frequency spectrum of 1300 Europeans from the Exome Sequencing Project ESP6400 dataset, 1298 Danes from the LuCamp dataset, and 432 Europeans from the 1000 Genomes Project to estimate the DFE of deleterious nonsynonymous mutations. We infer significantly fewer (0.38-0.84x) strongly deleterious mutations with selection coefficient |s| > 0.01 and more (1.24-1.43x) weakly deleterious mutations with selection coefficient |s| < 0.001 compared to previous estimates. Furthermore, a DFE that is a mixture distribution of a point mass at neutrality plus a gamma distribution fits best to two of the three datasets. Our results suggest that nearly neutral forces play a larger role in human evolution than previously thought.

## INTRODUCTION

A fundamental concept in population genetics is the distribution of fitness effects (DFE). The DFE refers to the proportion of new mutations that have particular effects on fitness. The DFE is a crucial quantity in evolutionary genetics because it determines how selection affects genetic variation (Eyre-Walker and Keightley 2007), the conditions under which recombination could evolve (Keightley and Otto 2006), and the spectrum of mutations potentially involved in genetic diseases (Eyre-Walker 2010). Further, an accurate DFE is required for robust inference of the amount of adaptive evolution across taxa (Boyko et al. 2008; Gossmann et al. 2012; Castellano et al. 2016; Galtier 2016), a topic of widespread interest. Because this distribution is so important, considerable effort has been put forth toward estimating it in several species, including humans (Eyre-Walker et al. 2006; Keightley & Eyre-Walker, 2007; Boyko et al., 2008; Li et al., 2010), *Drosophila* (Keightley and Eyre-Walker 2007; Kousathanas and Keightley 2013), yeast (Koufopanou et al. 2015), orangutans (Ma et al. 2013), gorillas (McManus et al. 2015), and mice (Halligan et al. 2013). Many of these studies suggest that the DFE has a leptokurtic distribution, with a large proportion of nearly neutral mutations, as well as many strongly deleterious mutations. For example, previous studies in humans (Eyre-Walker et al. 2006; Boyko et al. 2008) have estimated the parameters of a gamma distribution for the DFE of new nonsynonymous mutations. These studies have found approximately 57-61% of new nonsynonymous mutations to be moderately to strongly deleterious (|*s*|>0.001), about 15-16% to be weakly deleterious (1*e*-4≤|*s*|<1*e*-3), and the remainder (24-28%) to be nearly neutral (**Figure 1**).

The DFEs estimated for humans by Eyre-Walker et al. (2006) and Boyko et al. (2008) have been widely used for human population genetics. For example, the DFEs from these studies have been used to estimate differences in the genetic load across human populations (Henn et al. 2016), to model the ancient introgression of Neanderthal alleles into humans(Harris and Nielsen 2016), as a model for the frequency spectrum of deleterious polymorphisms in simulating data for disease studies (Uricchio et al. 2016), to evaluate the contribution of background selection to diversity on the Y chromosome in humans (Sayres et al. 2014), and to estimate the strength of selection acting on disease genes (Moon and Akey 2016). While these previous studies of the DFE have had considerable impact in the field, it is important to appreciate that the estimates were made using small samples, either in terms of total individuals sequenced or in terms of the proportion of the total genome sequenced. As such, most of the variation segregating in those samples is likely to be neutral or nearly neutral. Inferences about the proportion of moderately and strongly deleterious mutations largely come from assuming the gamma distribution approximates the DFE well, and then tabulating the proportion in those categories predicted by the gamma distribution. In other words, these proportions were extrapolated, rather than observed directly from the data.

This extrapolation of the proportion of strongly deleterious mutations may not be accurate. A more recent study using exome sequencing data from 200 Danish individuals (Li et al. 2010) estimated a DFE that differs considerably from that inferred in Boyko et al. Specifically, Li et al. found a mixture distribution consisting of a neutral point mass and gamma distribution fit best to their data (**Figure 1**). Additionally, they estimated that only 1% of new mutations have |*s*| > 0.0001 (compared to 57% in Boyko et al.), and 78% of new mutations fall in the 0.0001 ≤ |*s*| < 0.001 range (compared to 15% in Boyko et al.). Li et al. attributed this difference in the DFEs to their study considering a larger sample of individuals. As such, they surmised that they sampled more weakly deleterious variants, allowing more accurate inferences. However, this explanation has not been tested using simulations or larger datasets. Thus, the proportion of moderately vs. strongly deleterious nonsynonymous mutations in humans remains elusive.

**Figure 1.**
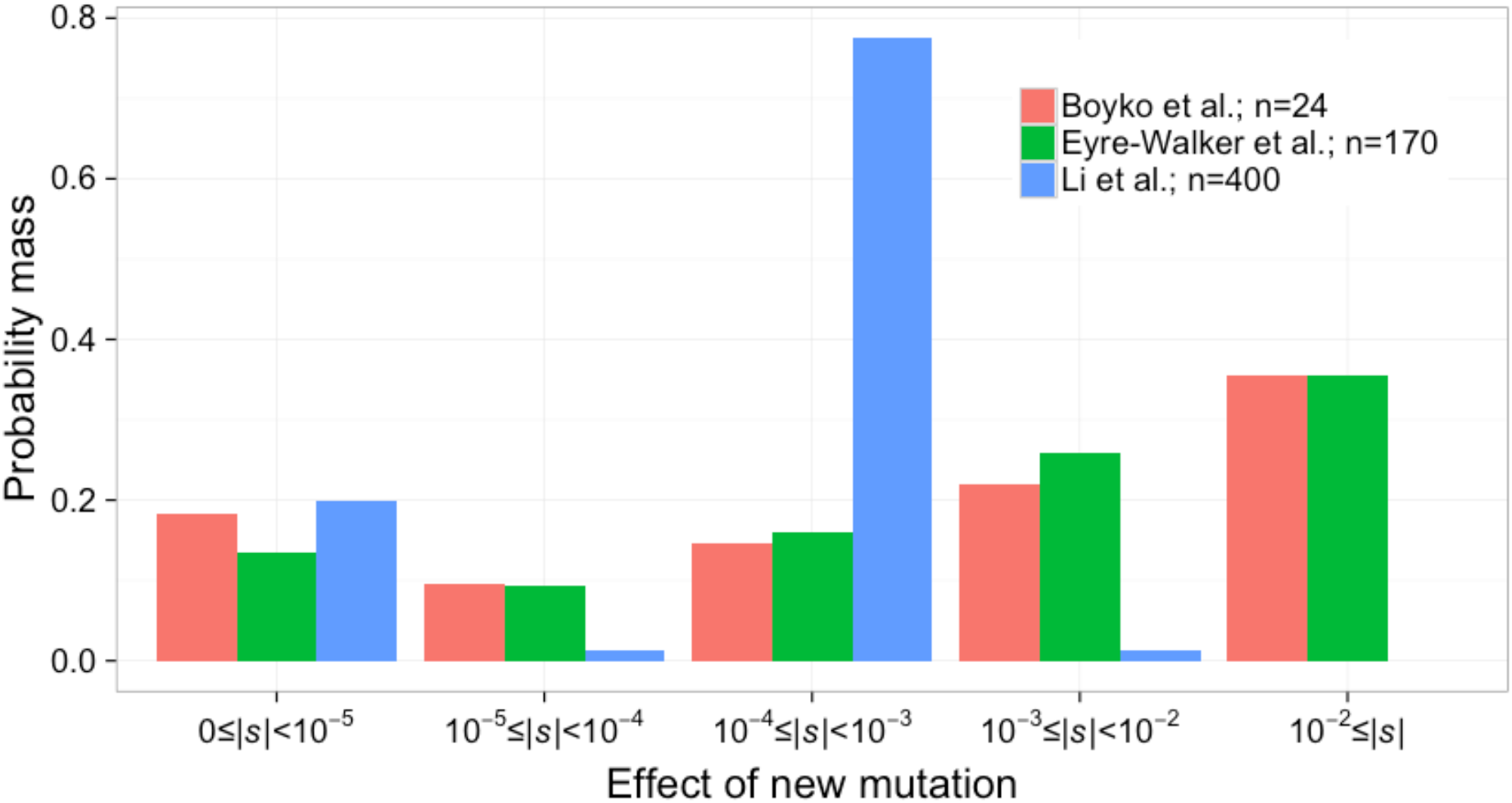
Previously inferred DFEs differ across studies. We rescaled the DFE in terms of the population size assumed or inferred in each study. A population size of 10,000 diploids is used to rescale the distribution of 2*Ns* to *s* for Eyre-Walker et al. (2006). For Boyko et al. (2008) and Li et al. (2010), we rescale the DFE from 2*Ns* to *s* using population sizes of 25,636 and 52,097 diploids, respectively (see Methods).

Due to large-scale genome and exome sequencing projects, samples of hundreds to thousands of individuals are available (Tennessen et al. 2012; Fu et al. 2013; Lohmueller et al. 2013; The 1000 Genomes Project Consortium 2015). These large datasets should yield more reliable inferences of the DFE because moderately deleterious polymorphisms should be segregating, albeit at low frequency, in these samples (**Figure S1**). As such, it should be possible to determine the functional form of the DFE and directly estimate the proportion of moderately and strongly deleterious mutations.

However, a major roadblock to using these new datasets for inference of the DFE is a lack of suitable software for inference from large samples. Generally, methods to infer the DFE summarize the genetic variation data of two classes of sites, one neutral and the other selected, by the site frequency spectrum (SFS). Then, they find the DFE that, under the inferred model of demography fit to the SFS from neutral sites, fits the observed SFS from selected sites. The method of Keightley and Eyre-Walker (2007), DFE-alpha, models demography using a Wright-Fisher transition matrix. It can only consider a demographic model with a single size change due to computational complexity. This is particularly limiting in large samples of human genetic variation since a single size change demographic model is insufficient for capturing the excess of rare variation in human populations (Keightley and Eyre-Walker 2007; Kousathanas and Keightley 2013). Another class of methods to infer the DFE uses the Poisson Random Field (PRF) approach (Sawyer and Hartl 1992; Hartl et al. 1994; Williamson et al. 2005; Eyre-Walker et al. 2006; Boyko et al. 2008; Li et al. 2010). This approach has been implemented in the program PRFREQ (Boyko et al. 2008), but that implementation becomes numerically unstable when applied to samples larger than a few hundred individuals. The program ∂a∂i (Gutenkunst et al. 2009) uses a similar framework, but implementing a DFE is slow due to the way that the DFE is integrated. Thus, there is a need for new software tools to infer the DFE from large samples.

In this study, we first extend the program ∂a∂i to analyze arbitrary DFEs in a computationally efficient manner. We implement these features in a module for ∂a∂i, which we call Fit∂a∂i. We then use this approach to estimate the DFE of deleterious nonsynonymous mutations from multiple large human datasets. We consider several different functional forms for the DFE. We find that across the multiple datasets, a mixture distribution where a proportion of mutations are neutral and the remainder are gamma distributed fits best. Analysis of multiple datasets suggests there are fewer strongly deleterious mutations where |*s*| > 0.01 (0.38-0.84x) than previously estimated in Boyko et al (35%), regardless of the functional form of the DFE. Further, our results are not consistent with a model where 99% of new mutations have a selection coefficient weaker than 1*e*-3, as suggested by Li et al. Because we anticipate that our estimates of the DFE will be useful in subsequent simulation studies, we provide SFS_CODE (Hernandez 2008) and SLiM (Messer 2013) commands to simulate from these DFEs.

## RESULTS

### Fit∂a∂i: Software to infer the DFE

Here we present our new software, Fit∂a∂i, to infer distributions of selection coefficients under the PRF model using the site frequency spectrum (SFS). Fit∂a∂i is a module that extends the functionality of the Python package ∂a∂i (Gutenkunst et al. 2009). Specifically, ∂a∂i uses diffusion theory to compute the expected SFS for a set of demographic parameters and selection coefficients. Fit∂a∂i offers a substantial computational improvement over the existing implementation of ∂a∂i for models involving more than a single selection coefficient. Essentially, Fit∂a∂i computes SFSs for a range of selection coefficients and saves each SFS into an array. Then, subsequent integrations of the DFE are done using the array of pre-computed SFSs. This process results in a substantial improvement in computational speed compared to the existing implementation of ∂a∂i, which re-computes the SFS for each selection coefficient in each step of the optimization process. Fit∂a∂i also allows parallel computation of the SFS by utilizing multiple cores or a cluster. Importantly, Fit∂a∂i leverages the modular nature of ∂a∂i to allow the user to define any arbitrary demographic models and DFE, including DFEs that are complex mixture distributions. Lastly, we incorporated an optimization routine that allows for constrained optimization of complex mixture distributions (see **Methods**).

### Datasets and demographic inference

We downloaded SNP data for: 432 unrelated European (EUR) individuals from the 1000 Genomes Project Phase 3 release (The 1000 Genomes Project Consortium 2015); 2000 Danish individuals from the LuCamp project (Lohmueller et al. 2013); and 6503 individuals from the NHLBI ESP6500SI-V2 European American (EUR) dataset (Tennessen et al. 2012; Fu et al. 2013)(Fu et al. 2013). The LuCamp and ESP datasets were projected down to sample sizes of 1298 and 1300 diploids, respectively, after filtering problematic individuals and for computational tractability. From these data we assembled the folded synonymous and nonsynonymous site frequency spectra (See **Methods**). We then fit a demographic model to synonymous sites using ∂a∂i. Briefly, this demographic model incorporates an Out-of-Africa bottleneck, a recovery period, and recent exponential population growth (**Figures S2** and **S3**). This methodology was repeated with each dataset projected down to *n*=24 chromosomes. Our estimates of demography as well as the inferred population sizes are presented in **Tables S1** and **S2**. Predictably, the parameter describing the magnitude of recent population expansion is difficult to infer with the smaller datasets. Although the demographic model we infer is biased by linked selection, this step controls for these effects during selection inference (Boyko et al., 2008; Kousathanas & Keightley, 2013; Messer & Petrov, 2013; Unpublished data from Huber, CD, Kim, BY, Marsden, CD, Lohmueller, KE, which was downloaded from http://biorxiv.org/content/early/2016/08/23/071209).

### Performance on simulated data

We verified the performance of Fit∂a∂i by simulating 100 datasets of sample sizes 24 and 2596 chromosomes under the PRF model using a demographic model of constant size, a 2x instantaneous size change, and the demography inferred from the LuCamp data described above (see **Methods**; **Table S1**, **Figure S2**). Data were simulated using the best-fit DFE of Boyko et al. (2008), rescaled to have an ancestral population size of *N*=10085 diploids (shape=0.184, scale=3226). Fit∂a∂i is able to accurately infer the DFE from our simulated datasets (**Table 1**). Predictably, the variance of our estimates of the most deleterious portion of the DFE (|*s*|>0.01) is 5 to 6-fold greater for the small samples. However, for the samples of size *n*=2596, the variance in the estimates of this portion of the DFE is significantly reduced and overall estimates of the proportions of the DFE where |*s*|>1*e*-3 are accurate. Therefore, this sample size should allow us to accurately infer the most deleterious portions of the DFE.

**Table 1.** Inference from Fit∂a∂i on simulated datasets.

| demography | $n_{chr}$ | shape | scale | $0 \leq \vert s\vert < 1e-5$ | $1e-5 \leq \vert s\vert < 1e-4$ | $1e-4 \leq \vert s\vert < 1e-3$ | $1e-3 \leq \vert s\vert < 1e-2$ | $1e-2 \leq \vert s\vert $ |
| --- | --- | --- | --- | --- | --- | --- | --- | --- |
| True<sup>a</sup> | -- | 0.184 | 3266<sup>b</sup> | 0.182 | 0.096 | 0.146 | 0.219 | 0.357 |
| Constant<br>size | 2596 | 0.180±0.010 | 3712.2±980.2 | 0.186±0.009 | 0.095±0.002 | 0.144±0.006 | 0.213±0.013 | 0.363±0.016 |
|  | 24 | 0.185±0.028 | 3613.1±4196.7 | 0.182±0.017 | 0.097±0.009 | 0.148±0.023 | 0.221±0.043 | 0.353±0.060 |
| 2x<br>expansion | 2596 | 0.191±0.007 | 2606.0±410.7 | 0.178±0.008 | 0.098±0.002 | 0.152±0.004 | 0.230±0.009 | 0.341±0.010 |
|  | 24 | 0.187±0.023 | 3259.8±2612.6 | 0.181±0.016 | 0.097±0.008 | 0.149±0.019 | 0.223±0.036 | 0.350±0.050 |
| LuCamp | 2596 | 0.182±0.008 | 3411.9±558.5 | 0.184±0.008 | 0.096±0.001 | 0.145±0.004 | 0.216±0.008 | 0.358±0.009 |
|  | 24 | 0.186±0.027 | 3435.4±3249.9 | 0.182±0.017 | 0.097±0.009 | 0.148±0.022 | 0.222±0.042 | 0.351±0.060 |

NOTE: 95% intervals were estimated as $\pm 1.96SD$ of 100 replicates in each simulation set.
<sup>a</sup>Values show the true DFE used to simulate the data.
<sup>b</sup>Here the simulation was scaled to an ancestral population size of $N=10085$ diploids.

Because it is not certain that the DFE is truly gamma distributed, we simulated datasets with DFEs from other distributions. Again, we scaled these DFEs to an ancestral population size of 10085 diploids. We considered the mixture distribution of Li et al. (2010), which consists of 20% neutral and 80% gamma-distributed (shape=4, scale=2.148) selection coefficients (herein called the “neutral+gamma DFE”). We also considered a mixture distribution consisting of 5 discrete bins (herein called the “discrete DFE”) with breaks at |*s*|=[0,1*e*-5,1*e*-4,1*e*-3,1*e*-2,1]. We examined three weighting schemes for this distribution. First, we computed the probability mass in each bin from the mixture distribution of Li et al. Then, we computed the probability mass in each bin from a gamma distribution with parameters: shape=0.203, scale=1082.1, but where the mass in the |*s*|=[1*e*-2,1] bin was placed into the |*s*|=[1*e*-3,1*e*-2) bin, and the opposite case where the mass in the |*s*|=[1*e*-3,1*e*-2) bin was placed into the |*s*|=[1*e*-2,1] bin. This was done to evaluate whether we could distinguish between these DFEs using the discrete DFE, and to evaluate our ability to distinguish strongly deleterious variants from moderately deleterious variants in a large sample.

We find that if the true underlying DFE is distributed according to the Li et al. neutral+gamma distribution, the discrete DFE is able to estimate the true DFE, albeit with some limitations. For example, when the true distribution is the Li et al. distribution, our inference under the discrete DFE distribution correctly estimates a negligible fraction of moderately or strongly deleterious new mutations (|*s*| > 1*e*-3) and correctly infers a mode of weakly deleterious new mutations (1*e*-4 ≤ |*s*| < 1*e*-3). However, estimates of the proportion of nearly neutral and neutral new mutations (|*s*|<1*e*-4) are less accurate (**Figure 2A**). When we simulate with the discretized distribution of Li et al., our estimates of the proportions of the discrete DFE are unbiased (**Figure 2B**). Additionally, we can distinguish between DFEs with varying proportions of moderately and strongly deleterious new mutations (**Figures 2C** and **2D**). Although it is unlikely that the DFE of any natural population is discretized in this manner, these results show that the discrete DFE can help to approximate the general form of the underlying DFE, even if the true DFE is multi-modal. This mimics what would be done in practice where the precise functional form of the DFE is not known *a priori*. Therefore, fitting the discrete DFE should provide a general idea of the true DFE, especially if the true DFE is significantly multi-modal. Notably, the discrete distribution should be able to distinguish between strongly and moderately deleterious mutations at our sample size of 2596 chromosomes.

**Figure 2.**
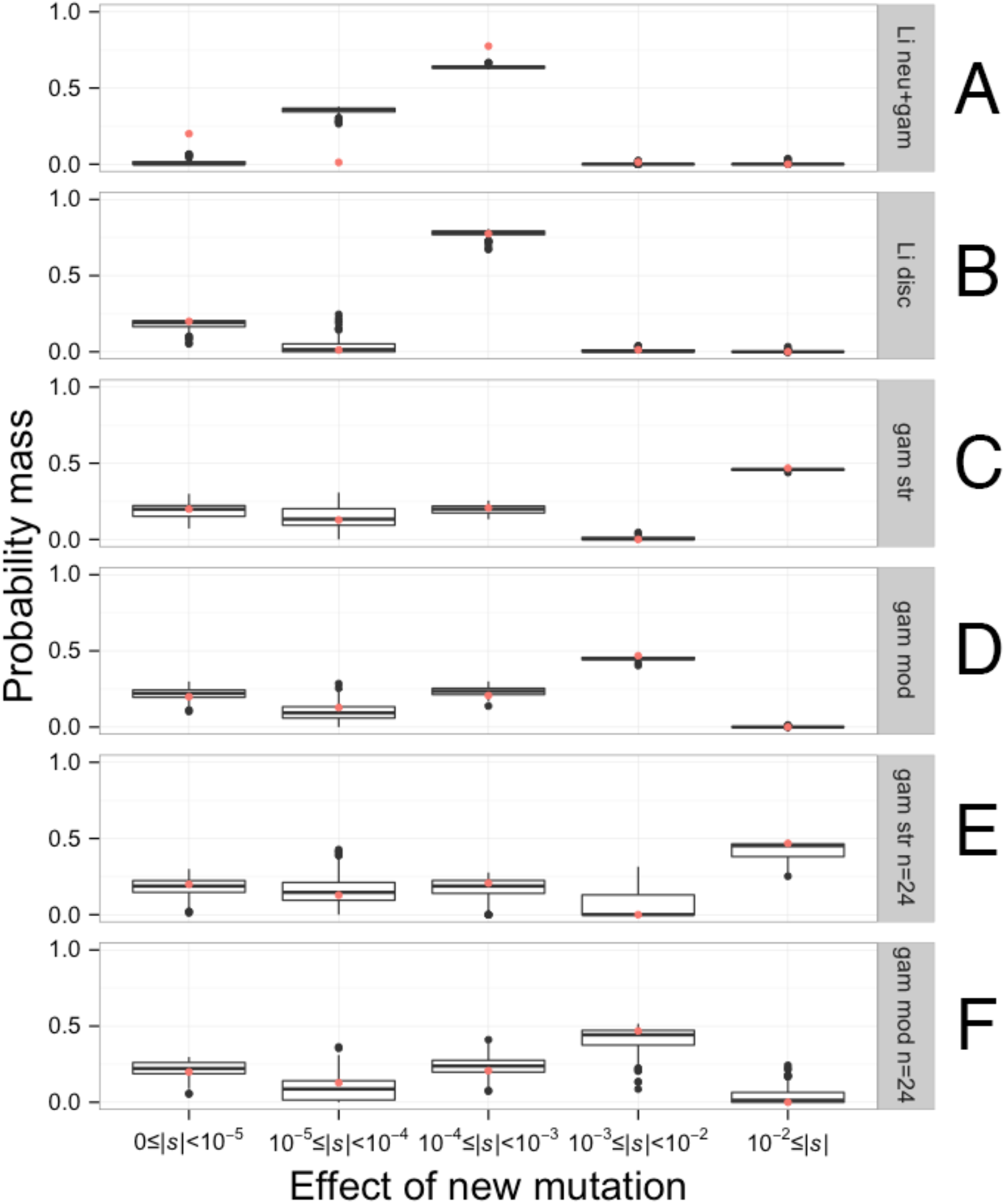
The discrete DFE can recover the approximate form of the DFE from simulated data. The distributions of the proportions of mutations with different selective effects, as inferred by the discrete DFE for 100 simulated datasets, are shown. Each simulation set assumed the demographic model fit to the LuCamp synonymous SFS. A red point depicts the true proportions of the simulated DFE. The true DFE for each set is: (A) the continuous neutral+gamma distribution of Li et al. (*p_neu_*=0.2, shape=4, scale=1.065*e*-4), (B) the discretized version of that distribution, (C-F) a gamma DFE (shape=0.215,scale=567.1), but where (C and E) the mass of the 1*e*-3≤|*s*|<1*e*-2 bin was added to the 1*e*-2≤|*s*| bin, and (D and F) where the mass of the 1*e*-2≤|*s*| bin was added to the 1*e*-3≤|*s*|<1*e*-2 bin. The datasets simulated for C and D had sample sizes of *n*=2596 chromosomes, while the datasets for E and F had sample sizes of *n*=24 chromosomes.

The procedure of first inferring demography from the synonymous SFS and then selection from the nonsynonymous SFS provides unbiased estimates of selection, even in the presence of linkage (Boyko et al. 2008; Messer and Petrov 2013; Comeron 2014). In other words, this methodology controls for the effects of selection at linked sites. However, it is unclear what effect population structure might have on inference of the DFE. It is well-known that such cryptic structure affects the SFS and may bias demographic inference (Ptak & Przeworski, 2002; Gazave et al., 2014). Further, large human resequencing datasets may contain cryptic population structure(Novembre and Ramachandran 2011). For example, the 1000 Genomes European sample is composed of 5 different subpopulations. To examine the performance of Fit∂a∂i when applied to datasets where the assumptions of the PRF model and a single, unstructured population are violated, we simulated 100 datasets with a linkage map, exon, and conserved non-coding regions based on the first 100Mb of chromosome 1 in SLiM (Messer 2013); Unpublished data from Huber, CD, Kim, BY, Marsden, CD, Lohmueller, KE, which was downloaded from http://biorxiv.org/content/early/2016/08/23/071209). Here, we simulated samples of size *n*=100 chromosomes sampled evenly across 8 genetically isolated subpopulations, each of size *N*=5,000 individuals, which all emerge from a single ancestral population of size *N*=5,000 individuals 400 generations ago (∼10,000 years). This population size increase reflects the Neolithic expansion into Europe under the demic diffusion model (Chikhi et al. 2002; Gazave et al. 2014). Additionally, we simulated selection on nonsynonymous sites following a gamma DFE with parameters: shape=0.2, scale=200.

Then, we fit a single population, single size change demographic model to synonymous sites, and a gamma DFE to nonsynonymous sites. This was done separately for each replicate. Again, we accurately infer selection from simulated data, even in the presence of linkage and population structure. Importantly, the single size change demographic model provides a reasonable approximation to both population expansion and structure (**Figure S4A**). This in turn allows for the accurate estimation of both the shape and scale parameters of the gamma distribution (**Figure S4B**). These results suggest that, provided that the demographic model fit to the data can adequately match the SFS of neutral synonymous sites, inferences of the DFE should be robust to cryptic, unmodeled, population structure.

### Validation of Fit∂a∂i by comparison to previous analyses

We also examined the performance of Fit∂a∂i on the unfolded African-American SFS from Boyko et al. (2008). Fit∂a∂i produces similar estimates of the shape and scale parameters of gamma distribution compared to Boyko et al. 2008 (Boyko: shape=0.184, scale=2488; Fit∂a∂i: shape=0.179, scale=3161). Additionally, Fit∂a∂i produced similar estimates of the proportions of mutations in different bins of the DFE (**Table S3**).

Therefore, simulations and a comparison to existing empirical data suggest that Fit∂a∂i can reliably infer the DFE in the presence of complex demographic scenarios. Below we present additional simulation scenarios below to examine the performance of Fit∂a∂i with varying sample sizes and when the assumed demography and DFE are mis-specified.

### Inference of the DFE from large datasets

Here we estimate the DFE for new nonsynonymous mutations using large samples. Specifically, we fit several distributions of the DFE to the folded nonsynonymous SFS of the 1000 Genomes EUR, Exome Sequencing Project EUR, and LuCamp datasets (**Table 2**). Unless otherwise noted, we assume a mutation rate of *μ*=1.5*e*-8 (Ségurel et al., 2014) and that the ratio of nonsynonymous to synonymous sites, *L*_*NS*_/*L*_*S*_, is equal to 2.31 (Unpublished data from Huber, CD, Kim, BY, Marsden, CD, Lohmueller, KE, which was downloaded from http://biorxiv.org/content/early/2016/08/23/071209). These values were used to obtain estimates of *N*, which were then used to convert estimates of 2*Ns* to estimates of *s* (see **Methods**). First, like previous studies, we fit a gamma distribution to the DFE (**Table S4**). We infer a strongly leptokurtic distribution, where there are many neutral and nearly neutral mutations, (i.e. 34-37% of new mutations have (|*s*|<1*e*-4), as well as a class of strongly deleterious mutations (i.e. 15-22% of new mutations have |*s*|>0.01). Interestingly, the estimates from the three different datasets are generally concordant, though the 95% confidence intervals (CIs) sometimes do not overlap. While this may suggest that the differences cannot be attributable to limited amounts of data in the SFS, we caution that these CIs are likely too narrow because they do not account for the non-independence of SNPs or the uncertainty of demographic estimates.

**Table 2.** Maximum likelihood estimates of various DFEs.

| Dataset | DFE | Parameter MLEs | Log-likelihood | AIC | $\vert \Delta\text{AIC}\vert ^b$ |
| --- | --- | --- | --- | --- | --- |
| LuCamp | gamma | shape=0.215,scale=562.1 | -3334.7 | 6673.4 | 13 |
| | neu+gamma | $p_{\text{neu}}=0.164,\text{shape}=0.338,\text{scale}=367.7$ | -3327.2 | 6660.4 | 0 |
| | neu+exp+let | $p_{\text{neu}}=0.304,p_{\text{exp}}=0.613,\text{rate}=66.56$ | -3337.8 | 6681.6 | 21.2 |
| | discrete<sup>a</sup> | $p_1=0.309,p_2=0.024,p_3=0.247,p_4=0.372$ | -3334.1 | 6676.2 | 15.8 |
| 1kG EUR | gamma | shape=0.186,scale=875.0 | -1450.5 | 2905.0 | 0 |
| | neu+gamma | $p_{\text{neu}}=0.031,\text{shape}=0.199,\text{scale}=820.6$ | -1450.8 | 2907.6 | 2.6 |
| | neu+exp+let | $p_{\text{neu}}=0.312,p_{\text{exp}}=0.509,\text{rate}=41.48$ | -1472.0 | 2950.0 | 45 |
| | discrete<sup>a</sup> | $p_1=0.286,p_2=0.099,p_3=0.222,p_4=0.305$ | -1453.4 | 2914.8 | 9.8 |
| ESP EUR | gamma | shape=0.169,scale=1327.4 | -3012.6 | 6029.2 | 2.6 |
| | neu+gamma | $p_{\text{neu}}=0.092,\text{shape}=0.207,\text{scale}=1082.3$ | -3010.3 | 6026.6 | 0 |
| | neu+exp+let | $p_{\text{neu}}=0.341,p_{\text{exp}}=0.504,\text{rate}=63.90$ | -3071.6 | 6149.2 | 122.6 |
| | discrete<sup>a</sup> | $p_1=0.334,p_2=0.041,p_3=0.201,p_4=0.306$ | -3029.5 | 6067.0 | 40.4 |

Note: These results are reported assuming $L_{NS}/L_S=2.31$ and $\mu=1.5e-8$ . See Table S4 for additional information.
<sup>a</sup>In terms of $|s|$ , these parameters correspond to the ranges of $|s|$ corresponding to: $[0,1e-5)$ , $[1e-5,1e-4)$ , $[1e-4,1e-3)$ , and $[1e-3,1e-2)$ , respectively. The mass in the $[1e-2,1]$ range is the complement of the total mass of the four aforementioned categories.
<sup>b</sup>Change in AIC relative to the model with the lowest AIC.

When compared directly to Boyko et al. (2008), the best-fit gamma DFEs inferred from all three datasets are generally shifted towards neutrality (**Tables 3** and **S4**, **Figure 3**). Here, we match the mutation rate of *μ*=1.8*e*-8 and *L_NS_*/*L_S_*=2.5 to provide a fair comparison(Boyko et al. 2008). This is because larger mutation rate assumptions should, given the same dataset, result in a DFE with more strongly deleterious new mutations. This occurs because fewer SNPs will be in the data relative to the mutation rate. Also note, we compare our results to what Boyko et al. observed in their African-American dataset, since the comparison is slightly more conservative than what was inferred in Europeans. However, the DFE fit to the European dataset in that study differs only by a few percent in each category of selective effect (**Table S5**). We infer 19.2%-22.9% of new mutations have selection coefficient |*s*|<1*e*-5, compared to the 18.3% observed by Boyko et al. (2008). This corresponds to 1.05x to 1.25x increases. Additionally, we infer 24.5%-29.8% of new mutations are strongly deleterious (|*s*|>0.01), which corresponds to 0.69x to 0.84x of the 35.5% inferred by Boyko et al. Taken together, when assuming a gamma distribution for the DFE, all three datasets suggest fewer strongly deleterious mutations than seen in Boyko et al, even when using the same mutation rate.

**Figure 3.**
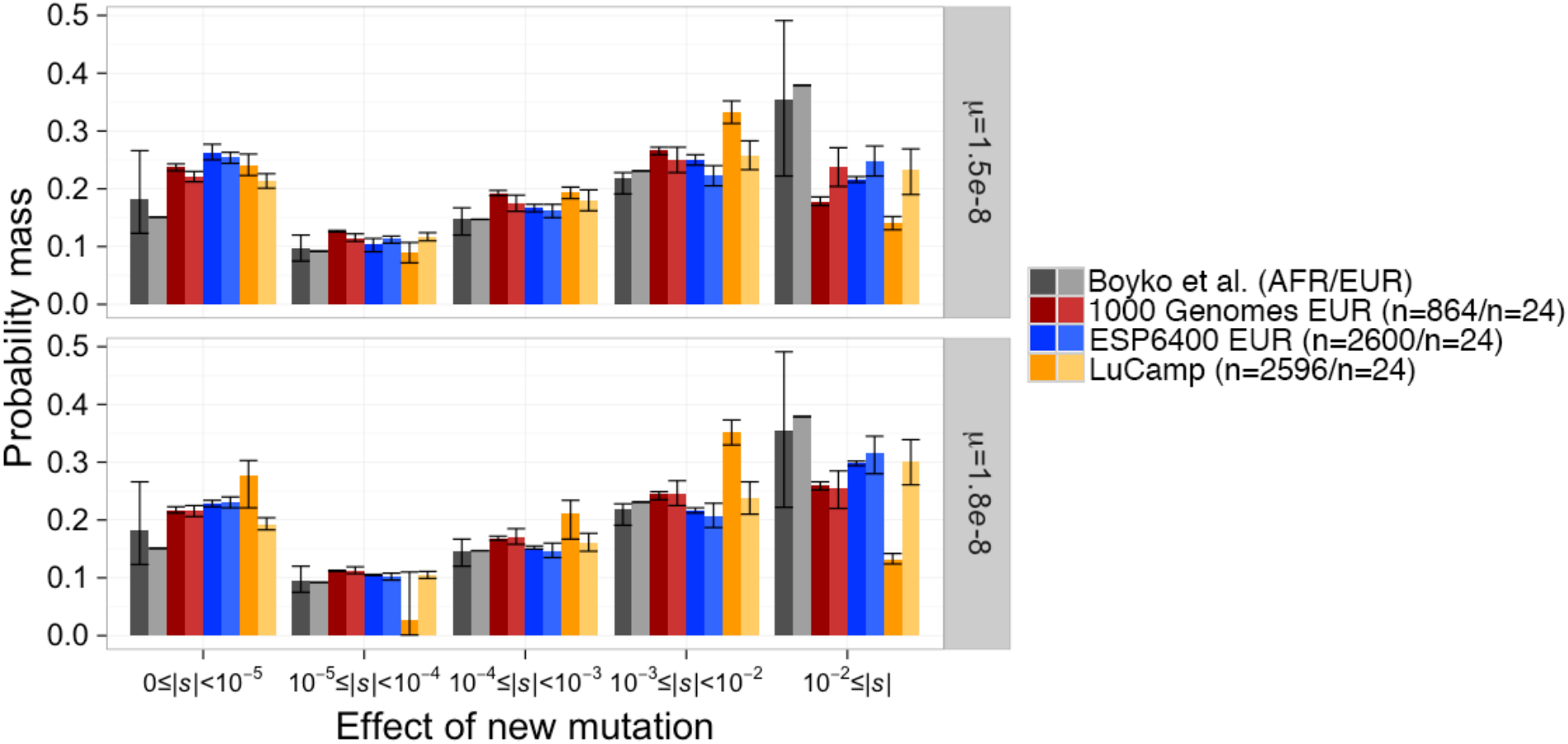
The distribution of selection coefficients of new mutations under our best-fit DFEs compared to Boyko et al. (2008). Results are presented for the best fit DFE to each full dataset and the best fit DFE when the data were projected down to *n*=24 chromosomes. Confidence intervals were estimated by Poisson resampling the nonsynonymous SFS and fitting a DFE 200 times. Confidence intervals for the DFE fit to the Boyko et al. European dataset were unavailable. Note that our models predict more nearly neutral mutations (0≤|*s*|<10^-5^) and fewer strongly deleterious mutations (0.01≤|*s*|) than Boyko et al., across all mutation rates. Top panel denotes our favored mutation rate while the bottom panel denotes the mutation rate used by Boyko et al. (2008).

**Table 3.**
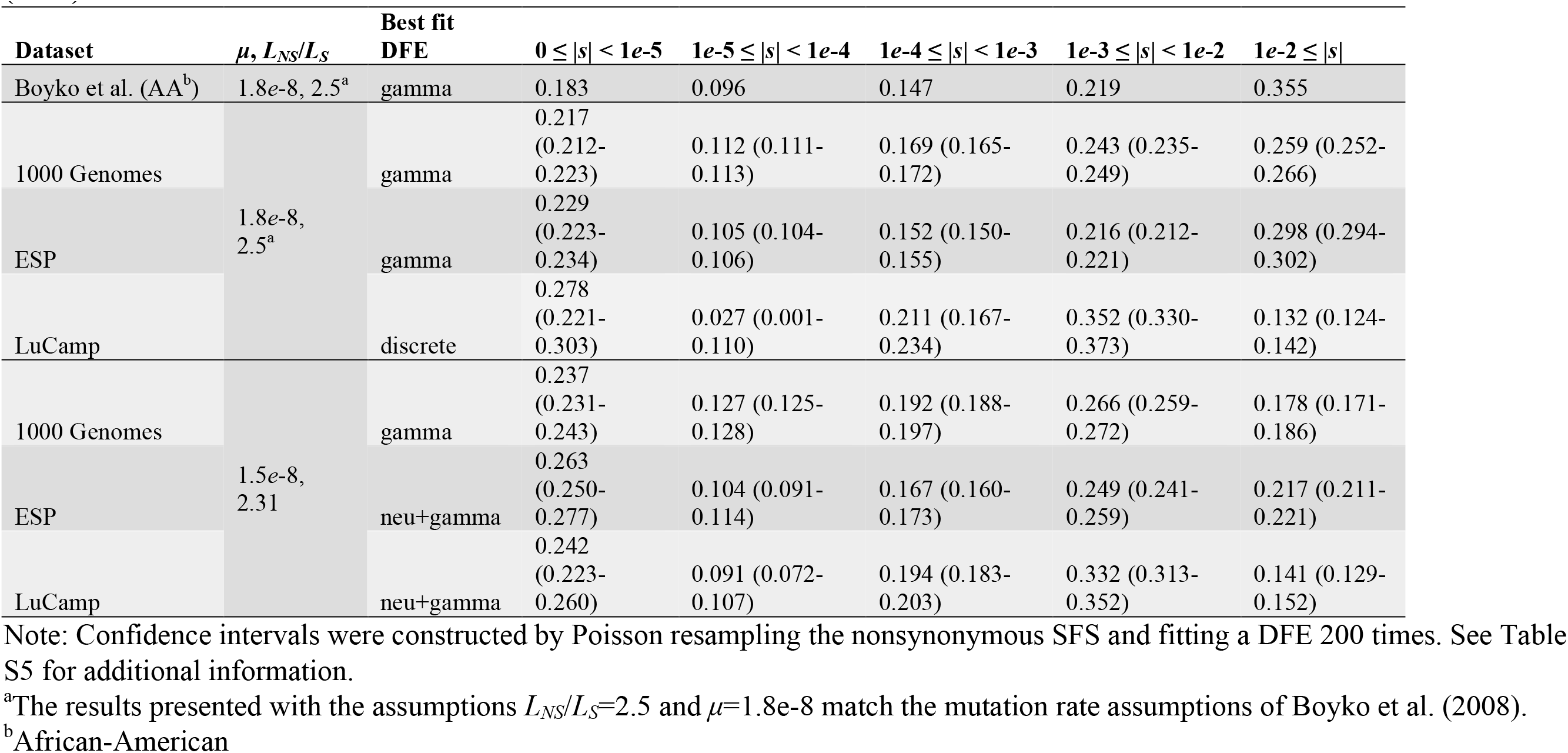
We infer more nearly neutral (|*s*|<1*e*-5) and fewer strongly deleterious (|*s*|≥0.01) new mutations than Boyko et al. (2008).

| Dataset | $\mu, L_{NS}/L_S$ | Best fit DFE | $0 \leq \vert s\vert < 1e-5$ | $1e-5 \leq \vert s\vert < 1e-4$ | $1e-4 \leq \vert s\vert < 1e-3$ | $1e-3 \leq \vert s\vert < 1e-2$ | $1e-2 \leq \vert s\vert $ |
| --- | --- | --- | --- | --- | --- | --- | --- |
| Boyko et al. (AA<sup>b</sup>) | $1.8e-8, 2.5^a$ | gamma | 0.183 | 0.096 | 0.147 | 0.219 | 0.355 |
| 1000 Genomes | $1.8e-8, 2.5^a$ | gamma | 0.217<br>(0.212-0.223) | 0.112 (0.111-0.113) | 0.169 (0.165-0.172) | 0.243 (0.235-0.249) | 0.259 (0.252-0.266) |
| ESP |  | gamma | 0.229<br>(0.223-0.234) | 0.105 (0.104-0.106) | 0.152 (0.150-0.155) | 0.216 (0.212-0.221) | 0.298 (0.294-0.302) |
| LuCamp |  | discrete | 0.278<br>(0.221-0.303) | 0.027 (0.001-0.110) | 0.211 (0.167-0.234) | 0.352 (0.330-0.373) | 0.132 (0.124-0.142) |
| 1000 Genomes | $1.5e-8, 2.31$ | gamma | 0.237<br>(0.231-0.243) | 0.127 (0.125-0.128) | 0.192 (0.188-0.197) | 0.266 (0.259-0.272) | 0.178 (0.171-0.186) |
| ESP |  | neu+gamma | 0.263<br>(0.250-0.277) | 0.104 (0.091-0.114) | 0.167 (0.160-0.173) | 0.249 (0.241-0.259) | 0.217 (0.211-0.221) |
| LuCamp |  | neu+gamma | 0.242<br>(0.223-0.260) | 0.091 (0.072-0.107) | 0.194 (0.183-0.203) | 0.332 (0.313-0.352) | 0.141 (0.129-0.152) |

Note: Confidence intervals were constructed by Poisson resampling the nonsynonymous SFS and fitting a DFE 200 times. See Table S5 for additional information.
<sup>a</sup>The results presented with the assumptions $L_{NS}/L_S=2.5$ and $\mu=1.8e-8$ match the mutation rate assumptions of Boyko et al. (2008).
<sup>b</sup>African-American

Next, we explored the fit of complex DFEs to these large samples. Using the same combination of mutation rates as with the gamma, we fit: the neutral+gamma mixture distribution; a mixture distribution of a point mass of neutral, a point mass of lethal, and exponentially distributed new mutations; and the discrete uniform DFE described previously. The MLEs are depicted in **Tables 2** and **S4**, as well as the proportion of mutations with varying selection coefficients predicted by these distributions (**Table S4**).

When we assume *μ*=1.5e-8 and *L*_*NS*_/*L*_*S*_=2.31, the neutral+gamma DFE fit best to the LuCamp and ESP datasets as reflected by the highest log-likelihood and AIC score (**Tables 3** and **S4**). The gamma still fit best to the 1000 Genomes dataset. Compared to the gamma DFEs inferred previously for two datasets, our best-fitting DFEs predict slightly fewer (0.92 to 0.98x) new mutations with |*s*|>1*e*-2, and slightly more (1.06 to 1.18x) new mutations of |*s*|<1e-5. When we matched the mutation rates of Boyko et al. with μ=1.8e-8 and *L*_*NS*_/*L*_*S*_=2.5, the discrete DFE fit best to the LuCamp dataset (**Tables 3** and **S4**). However, the gamma DFE continued to fit best to the 1000 Genomes and ESP datasets under these assumptions. Here, we predict significantly fewer (0.54x) new mutations have |*s*|<1*e*-2 compared to the gamma DFE for the LuCamp dataset.

The DFEs we have inferred thus far differ from that inferred in Boyko et al. (2008). In that study, 35.5% of new nonsynonymous mutations were inferred to be strongly deleterious in African-Americans, and 37.9% in Europeans. We infer fewer new strongly deleterious nonsynonymous mutations even when matching the mutation rates used in Boyko et al. Furthermore, our results remain consistent across datasets and assumed forms of the DFE.

Additionally, our estimates of the DFE differ substantially from the estimates provided by Li et al. (2010). Specifically, Li et al. infer almost no strongly or moderately deleterious new nonsynonymous mutations. That is, 1% of new nonsynonymous mutations have selection coefficients of 0.001<|*s*|<0.01 and 0% have selection coefficient |*s*|>0.01 (**Figure 1**). All of our estimates infer that at least ∼30% of new nonsynonymous mutations have selection coefficient |*s*|>0.001, even when the assumed mutation rate is small (**Table 2**, **Figure 4**). Our simulations suggest if the true underlying DFE follows that suggested by Li et al., we should at least be able to estimate those proportions using both the neutral+gamma and the discrete DFE (**Figures 2A** and **2B**). The fact that our inferences did not show similar estimates to those inferred by Li et al. suggests that our data and analyses are not consistent with the distribution inferred by Li et al. (**Table S5**). In the following sections, we explore several reasons why the different studies infer different DFEs.

**Figure 4.**
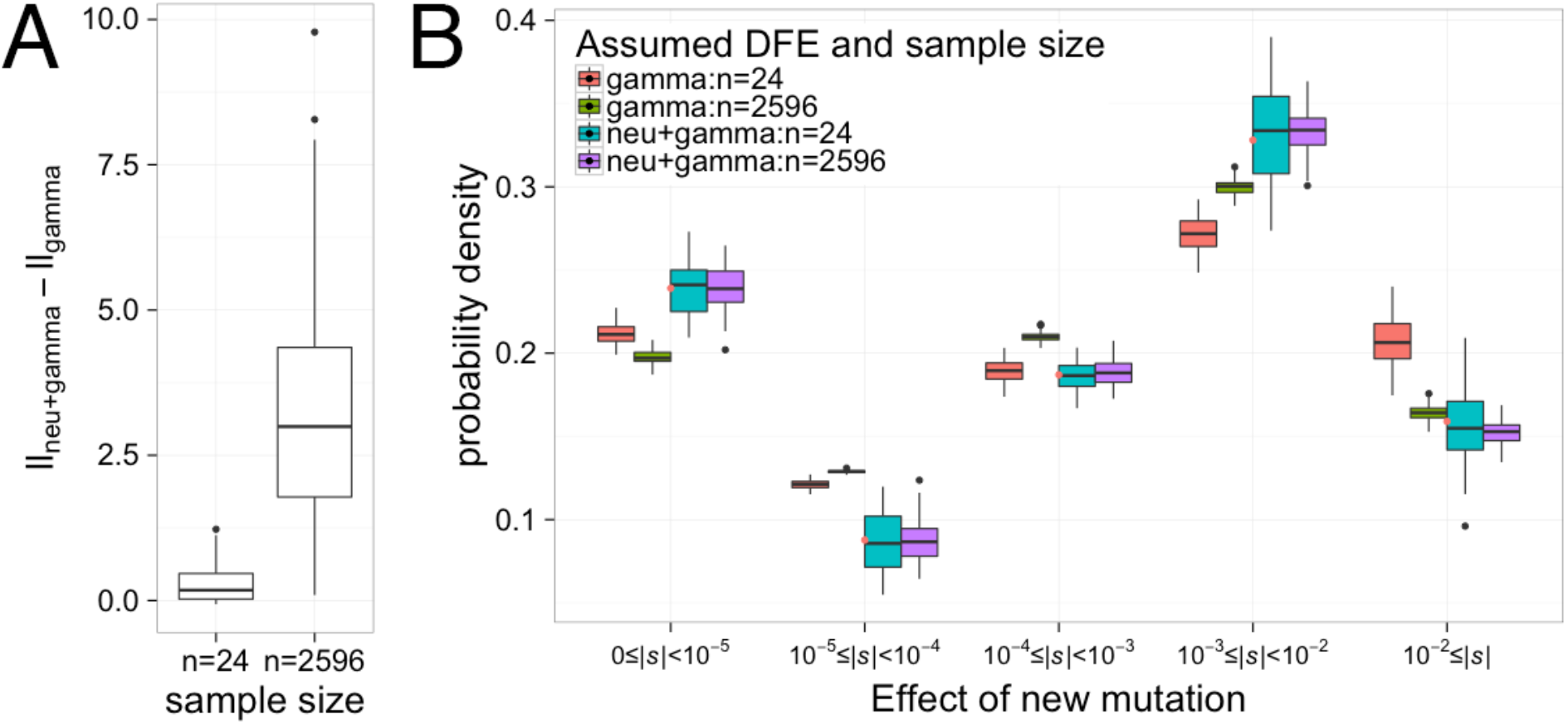
Small sample size and misspecification of the DFE can explain some of the differences between previous estimates and our estimates. Gamma and neutral+gamma DFEs were fit to 100 simulated datasets of sample sizes *n*=24 and *n*=2596 chromosomes where the true DFE was neutral+gamma distributed (*p_neu_*=0.164, shape=0.338, scale=358.8). (A) The distributions of the difference in log-likelihood between the gamma and neutral+gamma distributions. When the sample size is large (*n*=2596) the neutral+gamma distribution has a higher log-likelihood than the gamma distribution. However, the small samples (*n*=24) are unable to distinguish between the gamma and neutral+gamma distributions. (B) The estimated proportions of new mutations having different selective effects when fitting the gamma and neutral+gamma distributions. Note that when *n*=24, the gamma distribution over-predicts the proportion of strongly deleterious mutations (|*s*|≥0.01). Red dots denote the true proportion of mutations in each bin.

### Assessing the role of sample size using simulations

One possibility for the distinct estimates of the DFE is that the three studies used different sample sizes. Larger samples will have more moderately and strongly deleterious variants segregating than will smaller datasets (**Figure S1**). This could conceivably explain the different estimates. To investigate the effect of sample size on our ability to infer the DFE, we simulated 200 datasets, without linkage, of sample sizes *n*=12, 24, 100, 250, and 500 chromosomes. Each dataset was simulated using the demographic model and gamma DFE inferred from the LuCamp dataset.

First, we simulated neutral synonymous sites and inferred the demographic parameters from each dataset. This was done in two ways. First, we estimated the parameters from the full demographic model that was used to generate the data (herein the “full” model). Second, we inferred the parameters in a demographic model with only three instantaneous size changes (herein the “three-epoch” model). This is meant to mimic the situation in Boyko et al. where the true demography of the European population was likely complex, but simpler three-epoch demographic models could accurately fit the synonymous SFS. Next, as done in our inference and in previous studies, we estimated the parameters of a gamma distribution for the DFE for nonsynonymous mutations, conditioning separately upon the two demographic models.

When the full demographic model was fit to the simulated datasets, we found the variance of our estimates, both of demography and selection, decreased as sample size increased (**Figure S6**). We were unable to correctly infer the magnitude of recent population growth with small sample sizes as suggested by previous work (Keinan and Clark 2012; Nelson et al. 2012; Tennessen et al. 2012; Fu et al. 2013). However, this did not affect selection inference as long as the demographic model fit reasonably well to synonymous sites (**Figure S6**). At small sample sizes, the three-epoch model was able to fit the synonymous SFS well and thus estimates of selection were also unbiased. However, we found that for sample sizes larger than 100 chromosomes the three-epoch model increasingly became unable to account for the excess of rare variants caused by recent growth. The inability to account for the rare variants in the sample then biased the estimates of both the shape and scale parameters of the gamma distribution. However, this effect seems to be negligible at a sample size of 24 chromosomes (**Figure S6**).

When the true DFE is gamma distributed, small sample sizes can estimate the DFE, even when using an incorrect demographic model as long as that model fits the observed SFS of synonymous sites. The accuracy of the estimates increases with sample size, especially for the scale parameter, and notably provides better estimates of the strongly deleterious portion of the DFE (**Table 1**, **Figure S6B**). Thus, the results of Boyko et al. are unlikely to be affected by misspecification of demography due to small sample size.

Another possibility for the varying estimates of the DFE is that the DFE itself may be misspecified. Although parametric distributions are convenient for approximating the DFE, the true form of the DFE is unknown. Additionally, we have shown that the neutral+gamma DFE and the discrete DFE can fit large datasets better than the gamma DFE. To investigate an example of what would happen if the DFE is misspecified, we simulated 100 datasets without linkage for the best-fit neutral+gamma DFE inferred from the LuCamp dataset, scaled to an ancestral population size of 10085 diploids (*p_neu_*=0.164, shape=0.338, scale=358.8). We also downsampled each dataset to *n*=24 chromosomes. Then, we fit a gamma and neutral+gamma DFE to each full and downsampled dataset.

When the true DFE is neutral+gamma distributed, inference of the DFE from small samples overestimates the proportion of strongly deleterious new mutations. When the DFE is correctly specified, we obtain unbiased estimates of the DFE even from the small sample (**Figure 4B**). However, at sample size *n*=24 chromosomes, both the gamma and neutral+gamma distributions fit similarly to the data in terms of log-likelihood (**Figure 4A**). This was also observed in Boyko et al. (2008). Then, the extra parameter in the neutral+gamma distribution penalizes the true DFE when choosing the best-fit DFE by AIC score. This leads one to choose the gamma distribution as the best-fit DFE to the small sample, even when the true DFE follows a neutral+gamma distribution. Fitting the gamma distribution additionally results in a prediction of more new mutations with |*s*|>0.01 and a decrease in the proportion of new mutations with |*s*|<1*e*-5 than the true DFE (**Figure 4B**).

### Assessing the role of sample size using real data

Next, we investigated the role that sample size has on inference of the DFE from real data. To do this, we projected our synonymous and nonsynonymous frequency spectra down to a sample size of *n*=24 chromosomes to match the sample size of Boyko et al., then fit a demographic model and gamma DFE as previously described. Here we used the mutation rate assumptions *μ*=1.5*e*-8 and *L_NS_*/*L_S_*=2.31, but also matched the mutation rate of Boyko et al. (*μ*=1.8e-8 and *L_NS_*/*L_S_*=2.5). Then, we fit the gamma, neutral+gamma, and discrete, the best-fitting distributions to the full data, to the downsampled datasets.

As predicted by our simulations, there is generally little difference in the fit of the different DFEs to the downsampled datasets, in terms of log-likelihood (**Table S5**). The neutral+gamma and discrete DFEs often fit better than the gamma, but the difference is log-likelihood is small (0.1-0.6 log-likelihood units). Thus, the gamma DFE is selected as the best-fit DFE for all downsampled datasets by AIC. These results mimic what was observed in our simulations, although the pattern is not wholly consistent across datasets and mutation rates. When we assume *μ*=1.8*e*-5 and *L*_*NS*_/*L*_*S*_=2.5, the gamma DFE fits best to both the full and downsampled 1000 Genomes and ESP datasets (**Figure 3** and **Table S5**). There also appears to be little difference between the gamma DFE fit to the full and downsampled data. To contrast, the discrete DFE fits best to the LuCamp data under these mutation rates. Additionally, the neutral+gamma fit best to the full ESP and LuCamp data when we assume *μ*=1.5*e*-5 and *L_NS_*/*L_S_*=2.31. The gamma DFE fit to the downsampled versions of these datasets predicts more strongly deleterious (|*s*|>0.01) and more nearly neutral (|*s*|<1*e*-5) new mutations (**Figure 3** and **Table S5**). The DFE fit to the 1000 Genomes data using the lower mutation rates does not follow this pattern. The gamma DFE fits best to both versions of the dataset, yet the estimates from the small dataset still predict more strongly deleterious new mutations. These results seem to corroborate the results from our simulation, that is, fitting a DFE using a small sample may result in misspecification of the DFE, which in turn may cause inaccuracies in the inferred proportions of the DFE. We believe this may explain some of the differences between the findings of Boyko et al. (2008) and the findings in this study.

### Assessing the role of the likelihood function

Next, we investigated the performance of the multinomial vs. Poisson likelihoods at inferring the DFE. In this study, as well as in Boyko et al., we fit the DFE using the Poisson likelihood, which uses an *a priori* estimate of the population-scaled mutation rate, *θ*, to fit the curvature of the SFS as well as the total number of SNPs. Too few segregating variants would suggest the presence of strongly deleterious variants that are not segregating in the sample (Boyko et al. 2008).

The multinomial likelihood fits the curvature of the SFS while conditioning on the total number of SNPs. As such, the number of SNPs provides no additional information. The multinomial inference is similar to how the DFE was inferred by Li et al. (2010) in that they only used information from the curvature of the SFS. Note, however, Li et al. fit the population frequency spectrum using a least-squares approach while the multinomial likelihood fits the sample frequency spectrum. As such, the multinomial likelihood function does not strictly mirror the procedure of Li et al. Using the multinomial likelihood is convenient because it does not require any prior assumptions about the population scaled mutation rate, θ, yet may be underpowered when trying to identify the proportion of strongly deleterious mutations, unless such variants are segregating in the sample.

To compare the two likelihood methods at varying sample sizes, we fit the full model to simulated datasets of *n*=12, 24, 50, 100, 150, 200, 250 chromosomes using both the multinomial and Poisson likelihoods (**Figure S7**). Again, we simulated 200 datasets at each sample size with the LuCamp demography and a gamma DFE with parameters: shape=0.203, scale=1082.1. In general, the accuracy of our shape parameter estimate improves as the sample size increases, and we find the multinomial and Poisson likelihoods can both be used to reasonably estimate the shape parameter. While this trend holds true for the scale parameter using the Poisson likelihood, we find that we are unable to accurately infer the scale parameter using the multinomial likelihood, even for a sample of n=250 chromosomes. For example, nearly 50% all of the parameter estimates lie close to the maximum bound and 25% lie close to the minimum bound allowed during optimization. We found that this result can be explained by the likelihood surface being exceptionally flat with respect to the scale parameter. In other words, we cannot estimate the strength of purifying selection using only the curvature of the SFS with these sample sizes. Therefore, because Li et al. fit only the curvature of the SFS and excluded rare variants (>2% minor allele frequency) in a sample of size *n*=400 chromosomes, the power to detect strongly deleterious variants may be quite low, resulting in different parameter estimates from those in Boyko et al. and our present work.

## DISCUSSION

We developed a computational method to infer the DFE from large datasets, then fit DFEs to the nonsynonymous SFS of 432 Europeans from the 1000 Genomes Project, 1300 Europeans from the Exome Sequencing Project, and 1298 Europeans from the LuCamp project. The new DFEs suggest there are fewer strongly deleterious mutations than previous estimates (**Figure 3**). Although we find a neutral+gamma mixture DFE fits best to the ESP and LuCamp datasets, a gamma DFE seems to be a better fit to the 1000 Genomes data (**Table 2**). Nevertheless, our best-fit DFEs predict a 0.38-0.84x decrease in estimates of the proportion of strongly deleterious new mutations compared to the current widely-used estimates of Boyko et al. (2008). Additionally, our estimates are robust to assumptions of the mutation rate or the assumed functional form of the DFE. We show small sample size can lead to incorrect estimates of the DFE, specifically when the DFE is approximated with a parametric distribution that is not the true distribution (**Figure 4**). Therefore, our estimates provide an important update to previous studies of the DFE that used smaller sample sizes. Our current estimates of the DFE should be more reliable and precise, particularly the estimates of the proportion of moderately and strongly deleterious mutations. To facilitate their utility for future researchers, we provide scripts for implementing these models on GitHub (see **Methods**).

Our results suggest misspecification of the DFE may explain some of the differences in the DFEs we infer from small and large datasets. This is particularly relevant because the true DFE is almost certainly not a parametric distribution. At small sample sizes, the DFEs we fit tended to fit similarly to the data, in terms of log-likelihood (**Table S4**). Therefore, the DFE that had fewer parameters (i.e. gamma) was selected as the best-fit DFE. Additionally, we infer more strongly deleterious (|*s*|>0.01) new mutations from the downsampled datasets. We showed through simulations that even if the true DFE is neutral+gamma distributed, a gamma DFE is selected as the best fit to a small sample. Furthermore, this leads to inaccuracy in recovering the true proportions of the DFE (**Figure 4**). While the neutral+gamma distribution is also unlikely to be the true DFE, our simulations reproduce the patterns observed when downsampling the real data. Therefore, large sample size does not only help to improve the precision of the estimated DFEs, but also helps to approximate the correct form of the DFE. We expect this question to be better resolved as additional and larger sequencing datasets continue to be generated in the future.

Additionally, our results show that estimates of the DFE are sensitive to assumptions of the mutation rate. For a given a dataset, assuming a higher nonsynonymous mutation rate will result in the inference of stronger purifying selection due to the increased number of SNPs expected (but not observed) relative to the nonsynonymous mutation rate. There are two assumptions that factor into the calculation of the nonsynonymous mutation rate. First, it is unclear what the true mutation rate is. Whole genome, pedigree-based estimates suggest a mutation rate of about 10^−8^ per base pair per generation, exome-based estimates suggest rates of 1.5x10^−8^, and phylogenetic estimates suggest a mutation rate in the range of 2.0-2.5x10^−8^ (Ségurel et al. 2014). Second, we infer a mutation rate from synonymous sites, but use that mutation rate to make an *a priori* assumption about the nonsynonymous mutation rate. Many studies have the nonsynonymous mutation rate at 2.5 times the synonymous mutation rate, but we believe 2.31 to be a more accurate estimate, as this takes into account the CpG mutational bias and a 3:1 transition:transversion ratio (Unpublished data from Huber, CD, Kim, BY, Marsden, CD, Lohmueller, KE, which was downloaded from http://biorxiv.org/content/early/2016/08/23/071209). These two factors combined can result in large differences in the DFE. For example, the gamma DFE fit to the LuCamp data predicts 15% of mutations to be strongly deleterious when assuming *θ*_*NS*_/*θ*_*S*_=2.31 and *μ*=1.5x10^−8^, but 25% of new mutations to be strongly deleterious when assuming *θ*_*NS*_/*θ*_*S*_=2.5 and *μ*=2.5x10^−8^. Although our results remain qualitatively consistent across the range of mutation rates, uncertainty about the true rate leads to uncertainty in estimating the DFE.

Another important aspect of our results is the consistency of our estimates of the DFE between datasets. Our estimates of the DFE suggest a skew towards neutrality compared to previous studies, and we infer a consistent range of neutral (|*s*|<1*e*-5, 24-26%), moderately deleterious (1*e*-3<|*s*|<1*e*-2, 25-33%), and strongly deleterious (14-22%) new mutations between the three datasets. The consistency of our results across datasets suggests our inferences are accurate and robust to differences in sampling from populations, sequencing, bioinformatic processing, and sample size. This suggests the DFEs we have inferred are reliable updates to the DFEs inferred by Eyre-Walker et al. (2006) and Boyko et al. (2008).

It is also worth noting that our methodology has key differences from that of Li et al. (2010). Li et al. estimated the DFE using the population frequency spectrum excluding rare variants (MAF<2%), under a constant size demographic model using a least squares method, and fit the curvature of the SFS while not considering the total number of SNPs in the sample. The extent to which these methodological differences as well as differences in sequencing or bioinformatic processing of the data between their study and our present study contribute to the different estimated DFEs remains unclear. However, we have shown for small and moderately sized samples, fitting only the curvature of the SFS is insufficient for estimating the scale parameter of the DFE. In other words, for smaller samples the number of SNPs in the data must be considered to estimate the proportions of moderately and strongly deleterious new mutations, since moderately to strongly deleterious mutations are unlikely to be found in the sample.

Although Fit∂a∂i was developed to work with large sequencing datasets, it still has several limitations. The inference framework we utilize becomes increasingly slower for larger samples and requires significant computational resources and time to initially generate the SFS for the range of selection coefficients. Additionally, the frequency spectrum becomes difficult to compute for larger selection coefficients (2*Ns*>10,000). This is mainly because finer integration grids must be used to accurately estimate low frequency variants. For example, integrating over the full range of possible selection coefficients for a species with an effective population size much larger than humans may be prohibitively slow, especially for sample sizes of thousands of individuals. Also, like the method of Boyko et al. (2008), our method assumes additive selective effects and should be interpreted as averaging of selection over all heterozygotes and genetic backgrounds. Nevertheless, we anticipate that our method will be useful for estimating the DFE across the tree of life as polymorphism datasets from different species continue to accumulate.

The inference of a DFE with fewer strongly deleterious mutations has important implications for medical genetic studies. Our results suggest that there may be more weakly and moderately deleterious nonsynonymous mutations than previously appreciated. These variants could possibly contribute to disease risk. However, these mutations could also confound statistical tests that compare observed levels of variation to those predicted by population genetic models. For example, using the DFE of Boyko et al. would predict less segregating deleterious genetic variation because more new mutations were estimated to be strongly deleterious and would not segregate in the sample. However, if those mutations were instead only moderately deleterious, some could drift up in frequency and actually segregate in the sample. Further, a common approach to modeling how deleterious variants affect complex traits (Eyre-Walker 2010) assigns mutational effects on a trait as a function of their effects on fitness. This approach has been widely used to quantify the architecture of complex traits (The DIAGRAM Consortium 2012; Mancuso et al. 2016), to study the effects of demography on traits (Lohmueller 2014a; Simons et al. 2014), and to assess the power of rare variant association tests (Uricchio et al. 2016). The accuracy and realism of these models depend on having accurate estimates of the DFE.

More broadly, our results have important implications for understanding and quantifying deleterious variants across human populations (Lohmueller et al. 2008; Simons et al. 2014; Lohmueller 2014b; Do et al. 2015). Specifically, the fate of strongly deleterious mutations is relatively insensitive to population demography. The fate of weakly and moderately deleterious mutations, however, is linked more closely with effective population size (Henn et al. 2016). Human evolution in particular is governed by nearly neutral processes due to relatively small effective population sizes. Then, a DFE containing fewer strongly deleterious new mutations suggests the nature of purifying selection in humans may be different from what is currently understood. For example, larger proportions of moderately and weakly deleterious mutations may suggest greater differences in the proportion of segregating deleterious mutations and genetic load between human populations (Henn et al. 2016). Accurate inferences of the DFE are critical in this regard as researchers begin to use explicit models of demography and selection to quantify differences in the amounts of deleterious variants across populations (Brandvain and Wright 2016; Gravel 2016).

## MATERIALS AND METHODS

### Inference

We inferred demography and selection from segregating sites in a maximum likelihood framework (Williamson et al. 2005, Boyko et al. 2008, Gutenkunst et al. 2009). To do this, we summarized synonymous and nonsynonymous sites with the site frequency spectrum. The SFS can be described as a vector, ***X*=[***X*_*1*_, *X*_*2*_,…,*X*_*n-1*_], in which each entry *X*_*i*_describes the number of SNPs at frequency *i* in a dataset of size *n* chromosomes. Alternately, the SFS can be described as the proportion of SNPs at frequency *i*, ***P*=[***P*_*1*_, *P*_*2*_,…,*P*_*n-1*_], where each entry is computed as *P*_*i*_=*X*_*i*_/(Σ_*i*_*X*_*i*_). In the Poisson Random Field framework, each entry in the SFS is assumed to be an independent Poisson random variable (Sawyer and Hartl 1992; Hartl et al. 1994).

A demographic model, the parameters of which are described by *ϴ*_*D*_, was fit to the SFS of synonymous sites with ∂a∂i (Gutenkunst et al. 2009). Here, ∂a∂i uses a diffusion approximation to compute the distribution of allele frequencies given some demographic model, *f*(*x*; *ϴ*_*D*_). Then, the expected number of SNPs at frequency *i* in a sample of size *n* chromosomes can be written as:

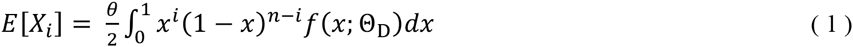

Then the multinomial likelihood is maximized to estimate the demographic parameters:

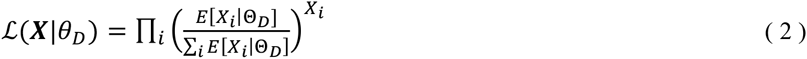

where *X*_*i*_ denotes the observed count of SNPs at frequency *i* in the sample. The multinomial likelihood is a convenient characterization of the SFS. Because it utilizes the proportions of SNPs at a particular frequency in the sample rather than the counts from the model, it does not require an *a priori* assumption of the mutation rate or ancestral population size. In other words, it fits the curvature of the SFS without considering the total number of SNPs in the data. The mutation rate of synonymous sites, denoted *θ*_*S*_, was then computed as the scaling factor difference between the optimized SFS and the data.

When fitting models incorporating periods of rapid exponential growth with ∂a∂i, we set the program parameter *dadi.Inference.set_timescale_factor*=1*e-*6, in contrast to the default setting of 1*e*-3. In ∂a∂i, periods of exponential growth are approximated with a series of instantaneous size changes, and if the step in time is not small enough, the recent growth parameters will not be inferred correctly. This causes the model to incorrectly infer the magnitude of recent growth, biasing inference of selection parameters. In other words, if there is an insufficient number of instantaneous size changes to approximate exponential growth, the number of predicted rare variants will be incorrect, which will in turn impact the selection inference.

### Selection inference

To infer the DFE, we developed fit∂a∂i, which utilizes ∂a∂i and some of the methodological improvements of Ragsdale et al. (2016). ∂a∂i is able to compute a distribution of allele frequencies *f*(*x*; *ϴ*_*D*_, *γ*), where *ϴ*_*D*_ is a vector containing the demographic parameters inferred from synonymous sites and *γ* is a single population-scaled selection coefficient. A DFE, denoted *g(*γ), can be incorporated by generating *f*(*x*; *ϴ*_*D*_, *γ*) for some range of *γ*, then weighting the contribution of each of these frequency spectra by *g(*γ) (Boyko et al. 2008).

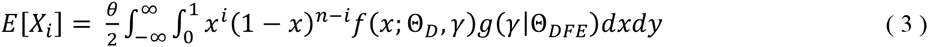

In the standard implementation of ∂a∂i, this process is time-consuming because the SFS must be recomputed repeatedly during each step of optimization. In other words, *f*(*x*; *ϴ*_*D*_, *γ*) is recomputed each time a given value of *γ* is evaluated in a DFE. This process can be especially slow for large ranges of *γ* and for large sample sizes. Therefore, similar to (Ragsdale et al. 2016), we initially computed the SFS for a range of selection coefficients, then cached these results to avoid re-computing the frequency spectra. In addition, we computed the many frequency spectra in parallel to save time; added compatibility for userdefined DFEs; modified the optimization routines available in ∂a∂i to work with cached spectra; and added the option to use constrained optimization for the inference of complex mixture distributions. These are the extensions that are part of the Fit∂a∂i module.

To infer selection, we fixed the demographic estimates to the MLEs of demography inferred from synonymous sites, 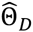. Then, we fit a DFE, the parameters of which are denoted as ϴ_DFE_, to nonsynonymous sites by maximizing the Poisson likelihood:

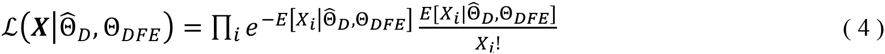

Unlike the multinomial likelihood, the Poisson likelihood requires an *a priori* assumption of the mutation rate of nonsynonymous sites, *θ*_*NS*_. To obtain this, we multiplied *θ*_*S*_ by an assumed ratio of nonsynonymous to synonymous sites, *L*_*NS*_*/L*_S_, to obtain *θ*_*NS*_. Each DFE is defined as an integrable function over a logspaced range of 600 selection coefficients over intervals between |*s*|=[1*e*-8,1]. We assumed any portion of the DFE smaller than |*s*|=1*e*-8 was effectively neutral (|*s*|=0), and any portion of the DFE where |*s*|>1 was lethal and would not be represented in our data. Note, here we only consider the deleterious DFE but the function can easily be extended to incorporate positive selection.

For mixture distributions incorporating a point mass at neutrality or lethality, we define the DFE so it can be treated as a single integrable function. We add the area of the point mass to a part of the distribution that is assumed to be neutral or lethal. For example, to add a point mass of neutral mutations to the neutral+gamma DFE, we add the probability mass of neutral mutations, *p_neu_*, to the |*s*|=[0,1*e*-8] portion of the distribution. Then, we integrate the gamma DFE between |*s*|=[0,1*e*-8] and sum it with p_neu_ to obtain the total mass of neutral mutations. Additionally, we utilized the SLSQP algorithm (Kraft 1988) as implemented in SciPy 0.17.1 to perform constrained optimization for mixture distributions incorporating more than two components.

### Estimating confidence intervals

To estimate confidence intervals for our data, we Poisson resampled the nonsynonymous SFS and re-fit the DFE to the resampled data (Boyko et al. 2008). We note our confidence intervals are probably too narrow for two reasons. First, we assume independence between all sites. Second, we do not account for the uncertainty in the demographic estimates since we condition upon the demographic model fit to synonymous sites.

### Estimating s from 2Ns

The DFEs inferred using the approach described above were for the population-scaled selection coefficient, γ =2*Ns*. Because we were interested in the distribution of *s*, we needed to estimate *N*. We computed *N* from the value of θ inferred from synonymous sites using the equation θ_S_=4*NμL*_*S*_ (**Table S2**). However, this value of *N* depends on assumptions of the mutation rate and the ratio of nonsynonymous to synonymous sites*, L*_*NS*_/*L*_*S*_, since these quantities are computed from the total number of coding sites, *L*_*S*_+*L*_*NS*_. Specifically, we assumed the mutation rate to be *μ*=1.5e-8 and *L*_*NS*_/*L*_*S*_=2.31, but also estimated the DFE assuming *μ*=1.8*e*-8 and *L*_*NS*_/*L*_*S*_=2.5*e*-8 to provide a fair comparison to Boyko et al. (2008), since the assumption of a higher mutation rate would suggest a DFE skewed towards more strongly deleterious new mutations. This is because there are fewer SNPs in the data relative to the nonsynonymous mutation rate. For the reanalysis of the Boyko et al. (2008) data, we assumed the same ancestral population size, *N*=7778 diploids.

To compute the total number of coding sites, *L*_*S*_+*L*_*NS*_, in each dataset, we intersected the coding exons from the UCSC canonical transcript with the relevant filters for each dataset. For the 1000 Genomes data, we intersected the Phase 3 strict mask, the exome targets, and the hg19 coding exons. For the analysis of the ESP (Tennessen et al., 2012; Fu et al., 2013) dataset, we used the intersection of the hg19 coding exons and the site-by-site annotations to count the total number of coding sites for which *n*≥2600 alleles had been sequenced. We obtained the value of *L*_*S*_+*L*_*NS*_ from Lohmueller et al. (2013). The quantities we used for these calculations are presented in **Table S2**.

### Simulations

To assess the performance of Fit∂a∂i, we performed forward-in-time simulations under different models of selection and demography. Simulations of independent sites were done using the program PReFerSIM (Ortega-Del Vecchyo et al. 2016), which simulates unlinked SNPs under the Poisson Random Field model. We simulated synonymous sites separately with a population-scaled mutation rate of *θ*=4000 to approximately match the amount of synonymous genetic diversity in our datasets. We simulated nonsynonymous sites at a ratio of 2.5 nonsynonymous to 1 synonymous site, in other words using *L*_*NS*_/*L*_*S*_=2.5. The demographic models and DFEs used were as described in the Results section. For simulations incorporating linkage and population structure we simulated 100 150Mb regions using the recombination rate and arrangement of functional elements on human chromosome 1 (Unpublished data from Huber, CD, Kim, BY, Marsden, CD, Lohmueller, KE, which was downloaded from http://biorxiv.org/content/early/2016/08/23/071209) using the forward simulation program SLiM (Messer 2013). We assumed a gamma DFE for both nonsynonymous (shape = 0.2, scale = 200) and conserved noncoding sites (shape = 0.0415, scale = 50; see Torgerson et al. 2009). We assume that 400 generations ago, the ancestral population expanded 8-fold and split into 8 genetically isolated populations. We then sampled 100 chromosomes equally across the 8 populations, combined them together, and analyzed them as though they were from a single population. The ancestral population was simulated for a burn-in period of 10*N* generations. To avoid prohibitively slow forward simulations, we simulated with an ancestral effective population size of 200 and scaled mutation rate, recombination rate, selection coefficients and demographic parameters accordingly (Aberer and Stamatakis 2013).

### Data

The 1000 Genomes Phase 3 data were downloaded from the European Bioinformatics Institute FTP site (http://ftp.1000genomes.ebi.ac.uk/vol1/ftp/release/20130502/). Related individuals were removed by sampling only mothers and fathers from trios or extended families. Only SNPs from the phase 3 exome targeted sequencing which passed the strict mask filter were used. The total length of sites considered in the analysis, that is, *L*_*S*_+*L*_*NS*_, was computed by taking this filtering into account. Variants were annotated using the 1000 Genomes Project filtered annotations. The ESP6400 dataset was downloaded from the Exome Variant Server (http://evs.gs.washington.edu/EVS/). Only sites with 1800 or more European individuals sequenced, according to the site-by-site annotations, were used for the analysis. This omission was factored into the estimation of *L*_*S*_ and *L*_*NS*_. The LuCamp data were obtained from Lohmueller et al. (2013). Detailed information about the total quantity of sites considered for our analysis can be found in **Table S2**.

### Software

This software is available at: https://github.com/LohmuellerLab/fitdadi.

## ACKNOWLEDGEMENTS

We thank Emilia Huerta Sanchez and Diego Ortega-Del Vecchyo for constructive comments on the manuscript. This work was supported by a Searle Scholars Fellowship and an Alfred P. Sloan Research Fellowship in Computational & Molecular Biology to KEL.

